# Rapid structural analysis of bacterial ribosomes *in situ*

**DOI:** 10.1101/2024.03.22.586148

**Authors:** Barrett M. Powell, Tyler S. Brant, Joseph H. Davis, Shyamal Mosalaganti

**Author notes:** Equal contributions (Joseph H. Davis, Shyamal Mosalaganti).

## Abstract

Rapid structural analysis of purified proteins and their complexes has become increasingly common thanks to key methodological advances in cryo-electron microscopy (cryo-EM) and associated data processing software packages. In contrast, analogous structural analysis in cells via cryo-electron tomography (cryo-ET) remains challenging due to critical technical bottlenecks, including low-throughput sample preparation and imaging, and laborious data processing methods. Here, we describe the development of a rapid *in situ* cryo-ET sample preparation and data analysis workflow that results in the routine determination of sub-nm resolution ribosomal structures. We apply this workflow to *E. coli*, producing a 5.8 Å structure of the 70S ribosome from cells in less than 10 days, and we expect this workflow will be widely applicable to related bacterial samples.

## INTRODUCTION

In the last decade, cryogenic focused ion beam milling (cryo-FIB) in conjunction with cryo-electron tomography (cryo-ET) has revolutionized structural biology^1-7^. Indeed, when combined with subtomogram averaging (STA), cryo-ET can produce structures at resolutions supporting molecular interpretations (∼3-10 Å), and it allows one to chart the dynamic landscapes of macromolecular complexes^8-11^. Critically, these combined techniques permit the characterization of such complexes directly in the native cellular environment (i.e., *in situ*), thereby enabling the discovery of novel (ultra-)structural states that may be lost during purification^6,12^. Further, cryo-ET uniquely reports the spatial distributions of macromolecular complexes relative to one another within the cell – often described as “molecular sociology,” which has led to key insights into cellular organization^13-16^. However, the practice of *in situ* cryo-ET is challenging. Among others, challenges include the slow production of lamellae, the number and quality of tilt series that can be acquired per lamella, and slow, labor-intensive data analysis that usually involves many rounds of computationally demanding sub-tomogram averaging, classification, and filtering^6^.

Several new approaches have been developed to address these aforementioned challenges. These include high-throughput sample preparation techniques, including the Waffle Method^17^ that permits the preparation of electron microscopy grids with high sample density; automated cryo-FIB milling with improved instrumentation to reduce ice deposition, which allows for longer milling times^18,19^; and parallel tilt-data acquisition with speeds of ∼5 min per tilt series^20-22^. Similarly, ever faster and more robust data processing software is continuously being developed^23-29^. Despite these advances, achieving high-resolution insights *in situ* typically remains a weeks-to-months-long challenge of navigating an array of sample preparation, data collection, and data processing strategies, even for large and highly abundant complexes such as ribosomes. Moreover, most cryo-ET campaigns aimed at sub-nm resolution structures have required the time-consuming acquisition of large datasets and, often, the development of bespoke dataset processing strategies (**Supplementary Figure 1**).

To help facilitate routine structural analysis *in situ* by cryo-ET, we developed a comprehensive sample preparation, imaging, and data processing workflow specifically designed to enable rapid structural analysis of abundant complexes, such as the ribosome in microbes. Here, we report applying this approach to the analysis of 70S ribosomes in *E. coli*. Our workflow produced a sub-6 Å resolution structure in less than 10 days when starting from cultured cells, and we expect this robust approach will be widely applicable for the high-throughput structural characterization of microbial ribosomes *in situ*.

## RESULTS

### A refined workflow for high-throughput data collection

Whereas some small (<300 nm thick) bacterial species are directly amenable to high-resolution cryo-ET, many species, including *E. coli*, are too thick (>500 nm) for imaging and thus must be thinned^30^. In this workflow, we used cryo-FIB milling to generate sections (lamella) of *E. coli* suitable for high-resolution imaging. Briefly, following Lam & Villa^31^, we applied a concentrated bacterial culture to a Quantifoil grid, resulting in a “carpet” of bacteria on the grid surface (**Figure 1a, b**; **Methods**). The thickness of these *E. coli* carpets was variable, ranging from 1-3 μm; therefore, to maximize throughput, we used milling angles of 6°-8° (corresponding to a stage tilt of 13°-15°), which allowed us to capture slices of multiple cells per lamella (**Figure 1b**). We typically obtained ∼20 lamella per 7-hour milling session (**Supplementary Figure 2**). After transferring these grids to a Titan Krios G4i transmission electron microscope, we performed automated data collection through SerialEM^32^, which allowed for microscope alignment, targeting and acquisition of 53 dose-symmetric tilt series per grid at an average rate of one tilt-series every 13 minutes. In sum, we produced a complete cryo-ET dataset within two days of harvesting our culture.

**Figure 1.**
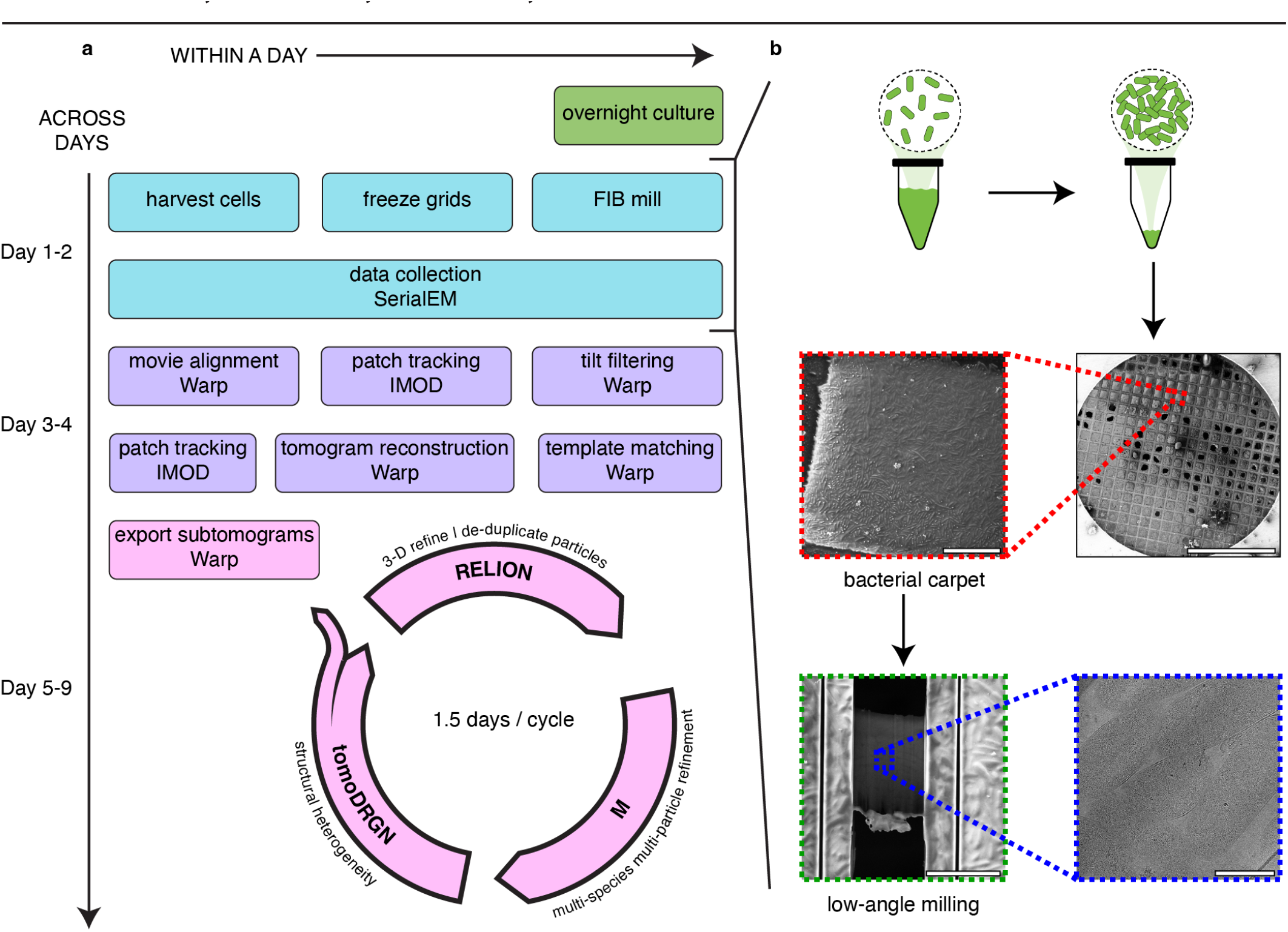
Modular workflow schematic for rapid structural characterization of microbial ribosomes *in situ*. **(a)** Sample preparation and data collection (cyan). Dataset pre-processing from movies to tomogram reconstruction (purple). Iterative sub-tomogram averaging and heterogeneity analysis (pink). **(b)** Representative scanning electron microscopy (SEM) overview image of the grid, SEM image of a single grid square (dotted red square), SEM image of a representative lamella from low-angle milling (dotted green square), and a low-magnification transmission electron microscopy (TEM) image of a lamella (dotted dark blue square). Scale bars, counterclockwise from top-right to bottom-right: 1 mm, 25 μm, 10 μm, 1 μm.

### A streamlined workflow for efficient data processing

We next employed a workflow to rapidly process our raw tilt-series data to high-resolution structures, specifically aiming to minimize time-consuming iterations (**Figure 1a**). Briefly, we performed movie alignment and micrograph CTF estimation in Warp^33^, followed by automated patch tracking in IMOD^34^. We excluded poorly aligning tilts in Warp and re-aligned the tilt-series in IMOD. Tilt series that remained poorly aligned were culled, resulting in 44 tilt series suitable for further analysis. Subsequently, we reconstructed highly binned tomograms in Warp for template matching and particle extraction, using a previously reported *E. coli* 70S ribosome structure obtained by subtomogram averaging (EMD-13270)^35^ as our template. We extracted 1,000 particles per tomogram, aiming to oversample the anticipated number of ribosomes by ∼3-fold. With this oversampling approach we expected to maximize the number of true positive ribosome picks at the expense of false positives, knowing we would subsequently filter particles using tomoDRGN, which we previously found to be highly efficacious at this task^28^. This data processing phase was completed within two days, resulting in a curated dataset of 44 tilt-series and 44,000 picked particles.

Extracted particles were progressively de-binned and subjected to 3-D auto-refinement in RELION version 3.1^23,36^, multiparticle coherent refinement in M^26^, and particle filtration in tomoDRGN (see **Methods**) over three cycles. Each cycle required ∼1.5 days of processing. Within each cycle, the initial RELION refinement against a lowpass-filtered reference improved each particle’s alignment, while subsequent M refinements improved per-tilt-series geometry and deformations, and tomoDRGN analysis facilitated efficient filtering of false positive particle picks. Re-extracting the filtered set of particles with finer pixel sampling at each cycle was designed to balance between sufficient image sampling to improve alignments and support particle filtering while enabling performant computation.

### Structural insights in situ within days

The first iteration of the RELION – tomoDRGN – M cycle with 5x binned particles allowed us to filter preliminary junk particles (**Figure 2a-c, Supplementary Figure 3a-c**), whereas the second iteration with 2x binned particles enabled clear assignment of remaining particles that constituted 70S ribosomes, 50S large ribosomal subunits, unassigned “junk” particles, and a minor population of false positives likely derived from platinum sputter (**Figure 2d-f, Supplementary Figure 3d-f**). For the third and final iteration, we skipped the RELION refinement due to minimal pose changes at this stage and instead directly used 1.5x binned particles for tomoDRGN – M analysis. This produced our highest resolution 70S ribosome structure at ∼5.8 Å (Nyquist limited to 4 Å resolution), with a local resolution distribution spanning 4-to-9Å (**Figure 2g, h**). In this map, we could resolve secondary structure elements throughout the ribosome and began to resolve bulky side chains in the high-resolution core (**Supplementary Figure 4**). In total, this data processing approach produced a “high” resolution consensus reconstruction, effectively filtered the particle stack to remove errantly picked particles, and primed the remaining particles for more detailed subsequent analysis within seven days of data collection.

**Figure 2.**
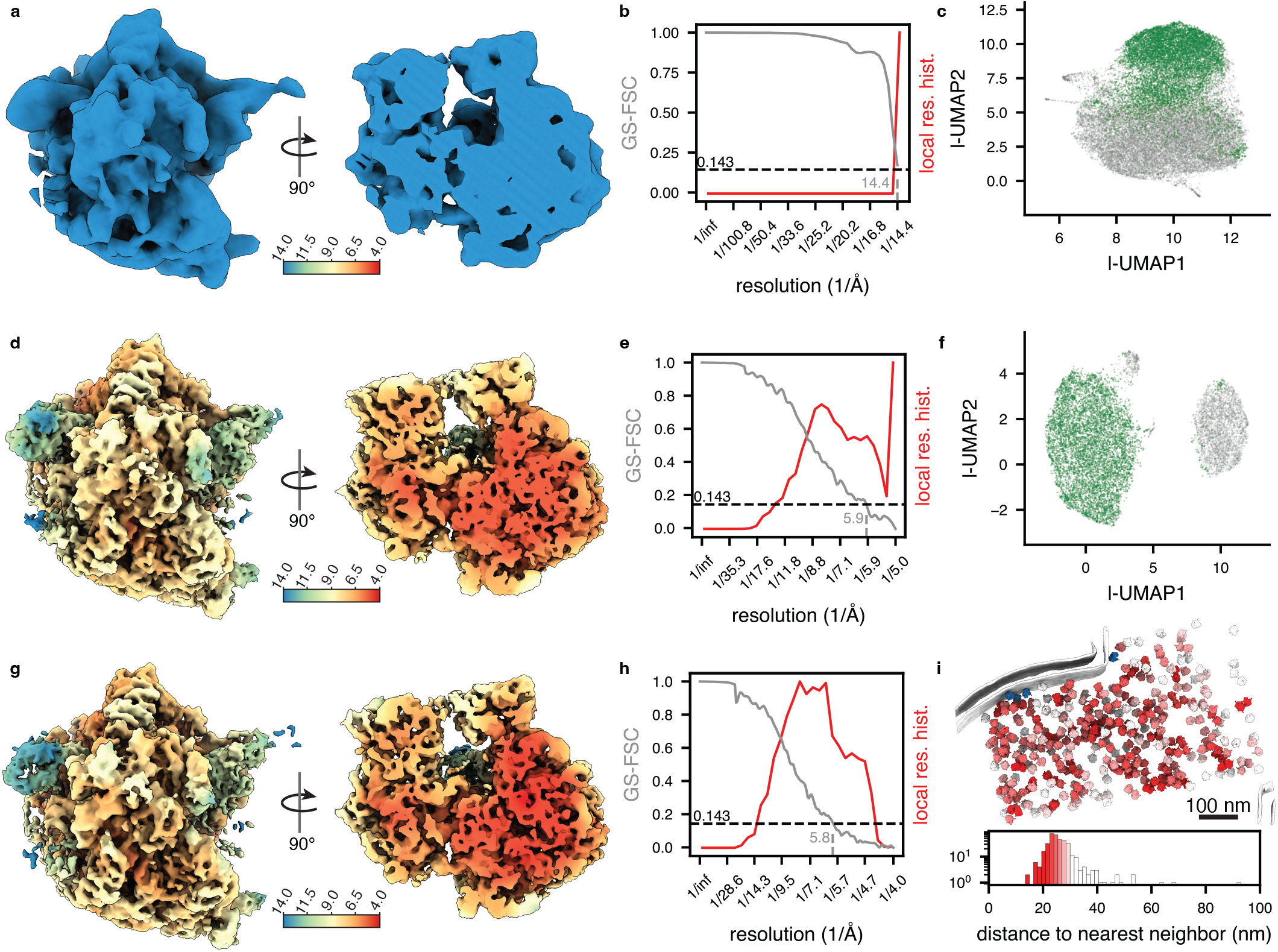
Iterative structural refinements enable rapid, high-resolution 70S ribosome reconstruction. **(a-c)** Data processing iteration 1 with 44,000 particles at ∼5x binning (7.03 Å/px). **(a)** Consensus subtomogram averaging (STA) reconstruction of bacterial 70S ribosome colored by local resolution (left) and cutaway view of the same (right). **(b)** Gold standard Fourier Shell Correlation (GS-FSC) from M (gray curve) with local resolution histogram overlaid (red curve). **(c)** TomoDRGN latent space visualized by UMAP dimensionality reduction, with particles selected for further processing indicated in green. **(d-f)** Data processing iteration 2 with 11,957 particles at ∼2x binning (2.5 Å/px). The panels are constructed consistent with the labels for (a-c). **(g-h)** Data processing iteration 3 with 8,170 particles at ∼1.5x binning (2 Å/px). Panels are constructed consistent with labels for (a-b). **(i)** Ribosome volumes from panel (g) mapped to corresponding cellular positions in the source tomogram (top). Cellular membranes (gray) were segmented with MemBrain v2^54^. Ribosomes are colored by the proximity to their nearest neighbor using a color scale depicted as a histogram (bottom). Membrane-associated ribosomes are colored in blue. Scale bar = 100 nm.

## DISCUSSION

Imaging macromolecules directly in their native cellular environment holds the great promise of interrogating the molecular sociology of the cell at molecularly interpretable resolutions. Moreover, by sidestepping the challenges associated with *in vitro* purification and reconstitution, it has already made great impact in guiding our understanding of the structure and function of critical macromolecular complexes^9,37-39^. However, routinely realizing the promise of *in situ* structural biology requires improved cryo-ET data collection and processing strategies. This becomes particularly apparent when considering the experimental and computational resources needed for cryo-ET compared to the highly efficient cryo-EM pipelines developed over the past decade^33,40-48^. Here, we report a rapid approach to structurally characterize bacterial ribosomes *in situ* that combines low-angle milling of bacterial carpets with automated tilt-series data collection and a streamlined data processing pipeline. We anticipate that this presented workflow will be directly applicable to the study of microbial ribosome biogenesis and translation and in structurally characterizing the impact of genetic, environmental, and pharmacological perturbations on the ribosome. Moreover, related workflows may prove helpful in analyzing other large, highly abundant complexes.

Obtaining thin, high-quality lamellae is a prerequisite for the structural characterization of protein complexes from thick cellular samples *in situ*. To increase the number of cells per lamellae, we plunged the bacteria at a concentration that resulted in ‘carpets’ on EM grids. We found that a trained operator performing low-angle, manual milling could produce ∼20 high-quality lamellae on a single grid in a 7-hour session (**Supplementary Figure 2**). We also show that a single day of automated imaging can produce a dataset suitable for relatively rapid structural characterization, with a sufficient number of particles to resolve secondary structural features. Finally, we found that the integration of tomoDRGN into our processing workflow expedited particle filtering, which is often rate-limiting in performing these analyses. We hypothesize that, like cryoDRGN^47,48^, tomoDRGN’s efficacy in this domain stems from its ability to project particles into a continuous multidimensional latent space that is highly effective in parsing the substantial heterogeneity between true positive particles and the many varieties of false positive particles that are often errantly picked from tomograms.

Whereas we have demonstrated the feasibility of our approach to resolve a ribosomal structure *in situ* with one week of processing, addressing specific biological questions will often require further specialized analyses. For example, to resolve, quantify, and spatially locate distinct functional states, one could leverage the generative capacity of tomoDRGN to reconstruct a large ensemble of heterogeneous structures to be queried on a per-particle basis for the occupancy of known structural elements with MAVEn^47,49^. Alternatively, automated (Kinman *et al*., in preparation) or expert-guided inspection of such an ensemble could enable the discovery of novel structural states lost through averaging in the consensus refinement. Importantly, we expect that the resolutions achieved using this workflow would allow the preliminary assignment of interacting macromolecules to unassigned density through the use of structural prediction tools such as AlphaFold^50-52^. Finally, we envision users performing tomogram-level analyses to examine spatial relationships between particles, membranes, filaments, or other cellular features identified in the tomogram (**Figure 2i**) using tools such as MemBrain, DeePiCt, NEMO-TOC, and others^53-56^.

We foresee ongoing improvements to this and related workflows, particularly those that either expedite the process or broaden the range of accessible macromolecular targets. For example, the rapid generation of the high-quality lamella is often limiting for cryo-ET, but the development of new instrumentation and control software that limits ice contamination, facilitates sample transfer to a cryo-TEM, and better automates the milling process is beginning to ameliorate this limitation^18,19,57^. For tilt-series data acquisition, we expect that machine learning approaches for automated lamella detection and target selection and the use of faster beam shifts rather than slower stage shifts during data acquisition will significantly expedite data collection^20,22^, potentially reducing the two days our workflow currently requires to collect datasets of sufficient size and quality for the analyses presented here. Additionally, recently developed software packages allowing for concurrent data collection and pre-processing would reduce our workflow by two additional days^58^. Separately, steady progress has been made in developing more accurate, efficient, and targetable particle-picking software, which should reduce the number of RELION – M – tomoDRGN cycles required to filter, align, and identify particles of interest^27,55,56,59^. Finally, ongoing work aims to develop unified processing workflows built for high-performance computing and on-the-fly data handling^24^, which could substantially reduce cycle time and improve attainable resolution.

## MATERIALS AND METHODS

### Bacterial culture

*E. oli* strain NCM3722^60^ was isolated from single colonies and grown in LB media without antibiotic selection at 37°C in a Multitron HT(Infors) shaker at 225 rpm to an OD_600_ of 0.5.

### Sample vitrification

Bacterial samples were pelleted at 500 x g for 5 minutes, supernatant was removed, and bacterial pellets were resuspended in LB to an OD_600_ of ∼20. Copper grids, 200 mesh, with holey R 1/4 carbon film (Quantifoil Micro Tools GmbH) were glow-discharged for 30 seconds at 5 mA using an EasiGlow system (Pelco) one hour prior to sample application. Grids were loaded into the environmental chamber of a Vitrobot Mark IV (ThermoFisher Scientific) set to 25 °C and 95% humidity. 3 μL of resuspended bacteria was applied to the carbon surface of the grid, and excess moisture was immediately blotted for 5 seconds at a blot force of 10. Whatman filter paper No. 1 was used to back-blot the grid, while a Teflon ring was placed on the sample-facing blot arm of the Vitrobot to simulate single-side blotting. Following blotting, grids were immediately vitrified by plunge-freezing in liquid ethane.

### Cryo-FIB milling

Plunge frozen grids containing bacterial samples were clipped into Cryo-FIB AutoGrids (ThermoFisher Scientific). Clipped grids were loaded onto the stage of an Aquilos 2 FIB-SEM (ThermoFisher Scientific), which was maintained at a temperature of ≤ -185°C. Grids were coated with a layer of organometallic platinum using a gas injection system (GIS) for 30 seconds and sputter-coated with inorganic platinum to form a protective/conductive surface for FIB milling. Tension-relief trenches were milled with a beam current of 1 nA. Subsequently, ∼5 μm thick lamella were milled at an angle of 6-8°. Following this, lamella were thinned to ∼1 μm thickness with beam currents of 100-500 pA. Milling progress and lamella integrity were checked via SEM imaging. Upon completion of all rough lamella on a grid, each was revisited and polished to a final thickness of ∼200 nm with the FIB at a beam current of 50 pA. All the lamella were polished within one hour post-rough-milling.

### Tilt series acquisition

Tilt series were acquired on a Titan Krios G4i microscope (ThermoFisher Scientific) operated at 300 kV and equipped with a Bioquantum energy filter (Gatan). Movies were collected at a magnification of 64,000x with a K3 Summit direct electron detector (Gatan) operating in super-resolution mode and binned 2x, corresponding to a nominal physical pixel size of 1.368 Å, with 8 frames per movie. Tilts were collected using SerialEM software^32^ with a modified dose-symmetric tilt scheme^61^ ranging from -54 to +66 degrees, starting from a +6° pretilt and at an interval of 3°. The total dose per tilt series was 100 e^-^/Å2. A 20 eV energy filter slit was used in the record mode, and the energy filter Zero Loss Peak was refined after each tilt series acquisition.

### Tomogram reconstruction

Raw tilt movies were aligned and initial CTF parameters were estimated in Warp (v1.1.0 beta)^33^. Unaligned tilt-series stacks were exported using tomostar mode in Warp. Tilt series (n=53) were aligned using in-house scripts (**Data and Software Availability**) that performed patch-based tilt alignment in IMOD^62^, sweeping over a range of patch sizes from 200 – 1,000 px at 4x binning, with each sweep iterated to prune poorly tracked patches. Tilts that were excessively dark, ice-contaminated, or poorly aligned in their respective tilt series were manually deselected in Warp before re-exporting unaligned tilt-series stacks for a second round of *de novo* tilt alignment. Tilt series with either weighted alignment residuals above 5 nm or fewer than 25 tilts were excluded from subsequent analysis. The remaining tilt series (n=44) were re-imported to Warp, specifying the IMOD-derived tilt alignment parameters and mdoc metadata filtered to exclude tilts that were de-selected in Warp previously. Improved CTF models per-tilt-series were fit in Warp, and tomograms were reconstructed in Warp at 10 Å/ px for inspection and particle picking.

### Particle picking

An *E. coli* 70S ribosome map (EMD-13270) was used for template matching after being Fourier cropped to 9.975 Å/ px, lowpass filtered to 40 Å, and real-space cropped to a 399 Å box in cryoSPARC^45^. Template matching was performed in Warp against 10 Å/px tomograms with 15° rotational steps and requiring particle separation of 50 Å center-to-center. The resulting particle picks were further filtered via in-house scripts (**Data and Software Availability**) to deduplicate particles within 75 Å (∼1/4 the diameter of a 70S) by pruning the lower figure of merit (FOM)-scoring particles. Finally, the top 1,000 particles, as judged by the FOM per tomogram, were used for subsequent analysis (n=44,000 particles).

### Sub-tomogram averaging and tomoDRGN analysis

In the first STA cycle, binned particles were extracted in Warp (pixel size = 7.03 Å/px, box size = 64px). An initial model for 3-D refinement was generated by CTF-corrected back-projection in RELION (v3.1)^23,63^ using the poses derived from template-matching. RELION 3-D auto-refinement with a 300 Å mask diameter was performed, attaining an unmasked resolution for the 70S ribosome of 15 Å with default parameters. Post-refinement global coordinates were used to filter duplicated particles within a cutoff distance of 150 Å, which removed 11,268 particles. The de-duplicated star file was downgraded to a RELION v3.0 star file using the relion_star_downgrade script^64^ and was used to extract particles as an “image-series” subtomogram in Warp at a pixel size of 7.03 Å/px and a box size of 64 pixels. A tomoDRGN model was trained on these particles for 50 epochs using an encoder A architecture 256x3, encoder B architecture 256x3, latent dimensionality 128, and decoder architecture 256x3. Manual inspection of 100 tomoDRGN-reconstructed volumes generated by *k*-means clustering of the latent space was used to exclude 20,819 particles we assigned as false positive / non-ribosomal (**Supplementary Figure 3**). TomoDRGN was used to filter the RELION star file, preserving the remaining 11,957 good particles for import into M (v1.0.9). M was used to perform multi-particle refinement using all tilts of each tilt series, a 300 Å particle diameter for the 70S ribosome, and using half-maps from RELION at a pixel size of 7Å. Four rounds of refinement were performed in M by iteratively adding new parameters to refine, which ultimately resulted in a Nyquist-limited 14Å resolution volume (**Supplemental Table 1**).

In the second STA cycle, particles were extracted at approximately 2x binning (pixel size = 3 Å/px, box size = 150px) from M. RELION was used for initial model generation, 3-D refinement to ∼9 Å GS-FSC resolution (unmasked) and particle de-duplication as described above. Particles were re-imported to M for four refinement cycles as described above, using half maps from RELION resampled to 2.5 Å/px. M refinement resulted in a 5.9 Å resolution map (**Supplemental Table 2**). Particles were exported as image-series subtomograms from M at a pixel size of 3 Å/px and a box size of 100px and used to train a new tomoDRGN model as described above but for modification of the decoder to a 512x3 architecture. As above, manual inspection of 100 tomoDRGN-reconstructed volumes from *k*-means clustering of the latent space was used to separate 70S ribosomes (n=8,170), 50S ribosomes (n=376), non-ribosomal particles (n=3,360), and strongly contrasting particles that mapped to platinum sputter (n=51).

In the third and final STA cycle, the 70S and 50S ribosomes were imported into M as distinct structural species, using half-maps from tomoDRGN image-series back-projection rescaled to 2 Å/px in RELION (approximately 1.5x binning). Only tilt series with at least 40 particles were refined. Twelve rounds of refinement were performed in M, ultimately reaching resolutions of 5.8 Å and 12.3 Å for the 70S and 50S, respectively (**Supplemental Table 3**).

## ACKNOWLEDGMENTS

We thank Dr. Vinson Lam from the University of Michigan Cryo-EM Facility (U-M Cryo-EM) for help with the data acquisition. U-M Cryo-EM is grateful for support from the University of Michigan Life Sciences Institute (LSI), the University of Michigan Biosciences Initiative, and the Arnold and Mabel Beckman Foundation grant. We thank the LSI-IT and the MIT-IBM Satori team for computing resources. We thank Laurel Kinman, Maria Carreira, and Chenxi Ou for helpful discussion and feedback. This work was supported by the NIH Director’s New Innovator Award (DP2) (1DP2GM150019-01) to SM; grants from the NIH (R01-GM144542) and NSF (CAREER-2046778), and awards from the Sloan Foundation and the MIT Jameel Clinic to JHD; and NIH grant 5T32-GM007287 to BMP. The funders had no role in study design, data collection and analysis, the decision to publish, or the preparation of the manuscript.

## CONTRIBUTIONS

JHD and SM designed the project. BMP and TSB prepared the samples and collected the cryo-FIB and cryo-ET data. BMP established the data processing workflow and performed all the analyses. BMP and TSB prepared the figures and wrote the first draft of the manuscript. All authors were involved in preparing the final draft of the manuscript.

## DATA AND SOFTWARE AVAILABILITY

The correspondence for material and data in the manuscript should be addressed to. The raw data described in this study has been deposited to EMPIAR under accession number EMPIAR-11983. The final consensus map has been deposited to the Electron Microscopy Data Bank (EMDB) under accession number EMD-44152. In-house scripts used for batch tilt series alignment and template matching particle filtration, as well as trained tomoDRGN models and analyses used during particle filtering, are available at https://doi.org/10.5281/zenodo.10841632.

## SUPPLEMENTARY INFORMATION

**Supplementary figure 1.**
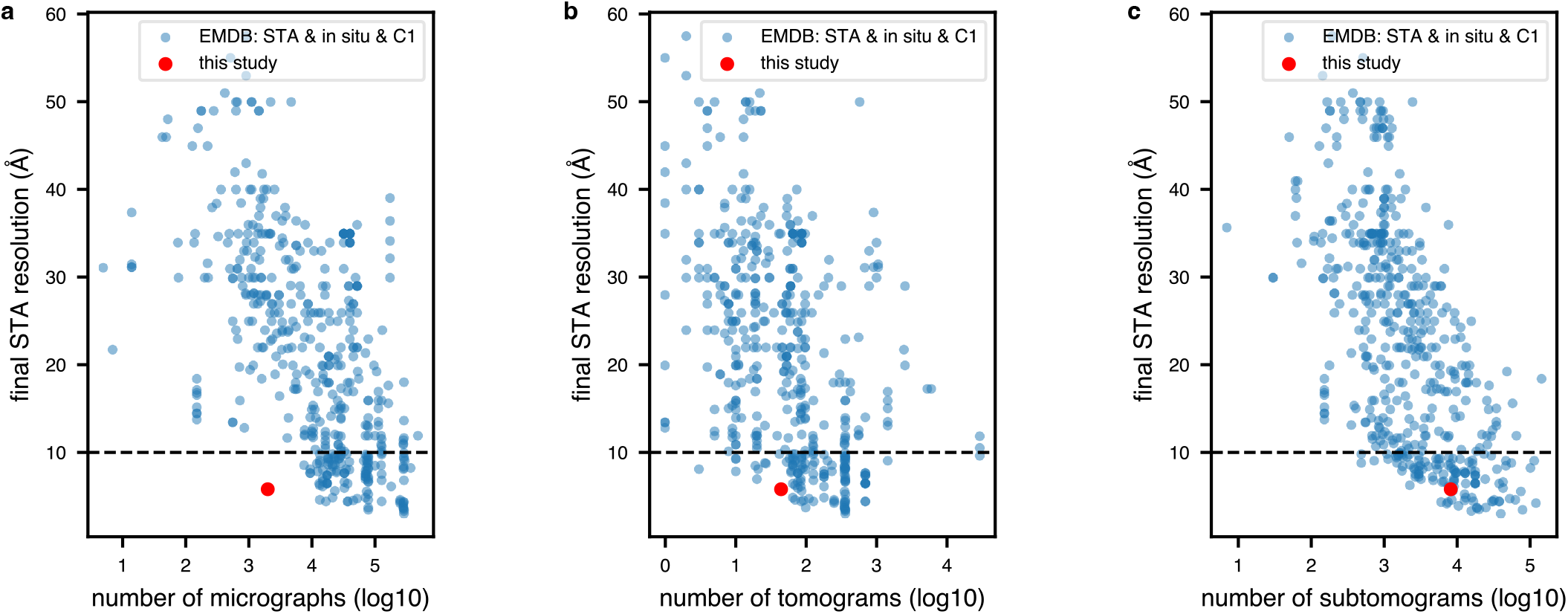
Select EMDB summary statistics for STA datasets and processing workflows. **(a)** Final STA resolution as a function of dataset size (by number of micrographs) for all EMDB entries through 15 March 2024, which are annotated as determined by STA with C1 point group symmetry from samples in the “cell” aggregation state. Note that this includes thinned/milled lamellae and samples directly imaged on grids. **(b)** Same as (a), but with dataset size plotted as number of tomograms. **(c)** Same as (a), but with dataset size plotted as number of subtomograms.

**Supplementary figure 2.**
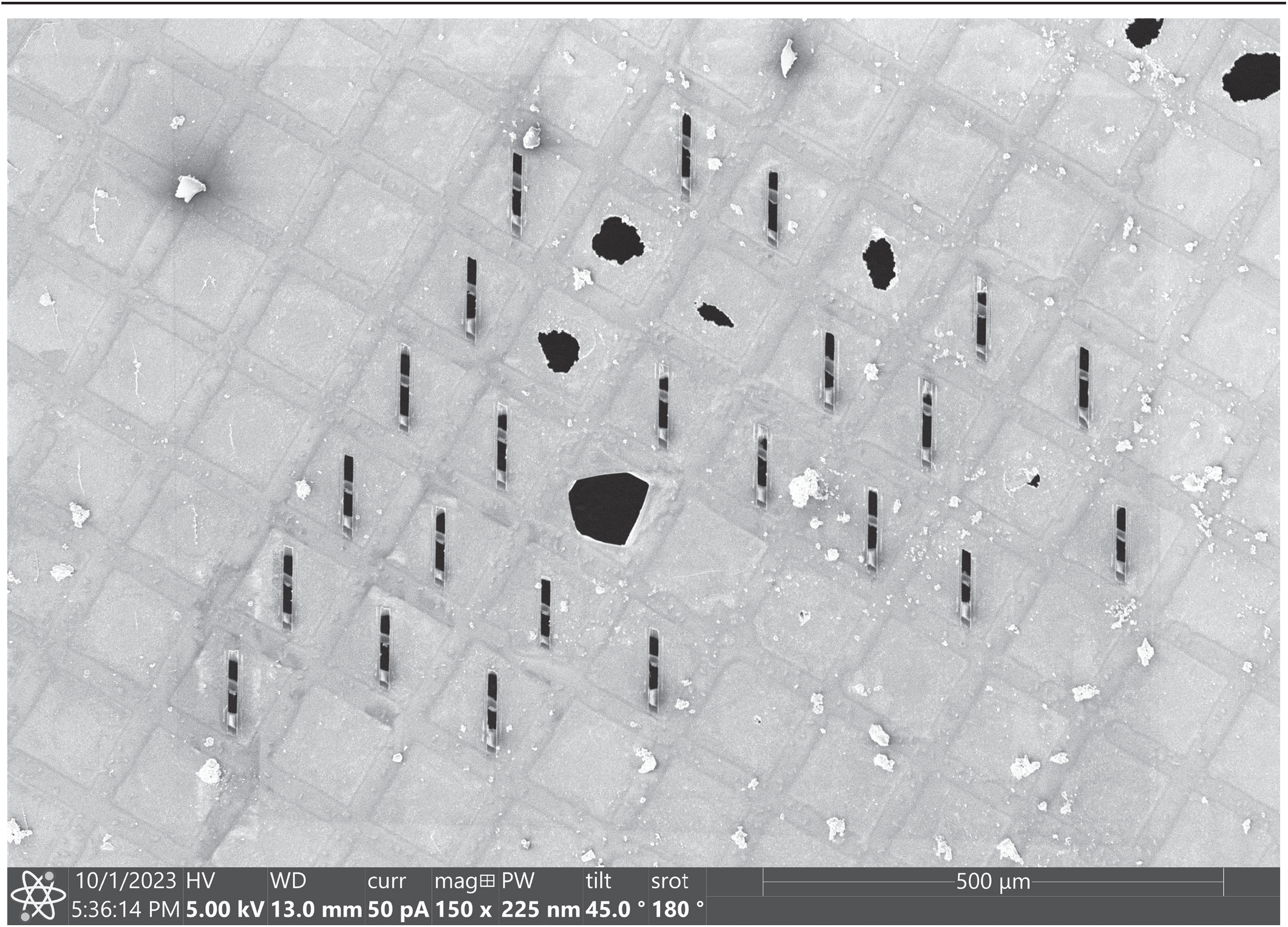
Cryo-focused ion beam (cryo-FIB) milling of *E. coli*. Representative scanning electron micrograph image of cryo-FIB milled *E. coli* grid showing 23, 10 μm-wide intact lamella. Scale bar: 500μm.

**Supplementary figure 3.**
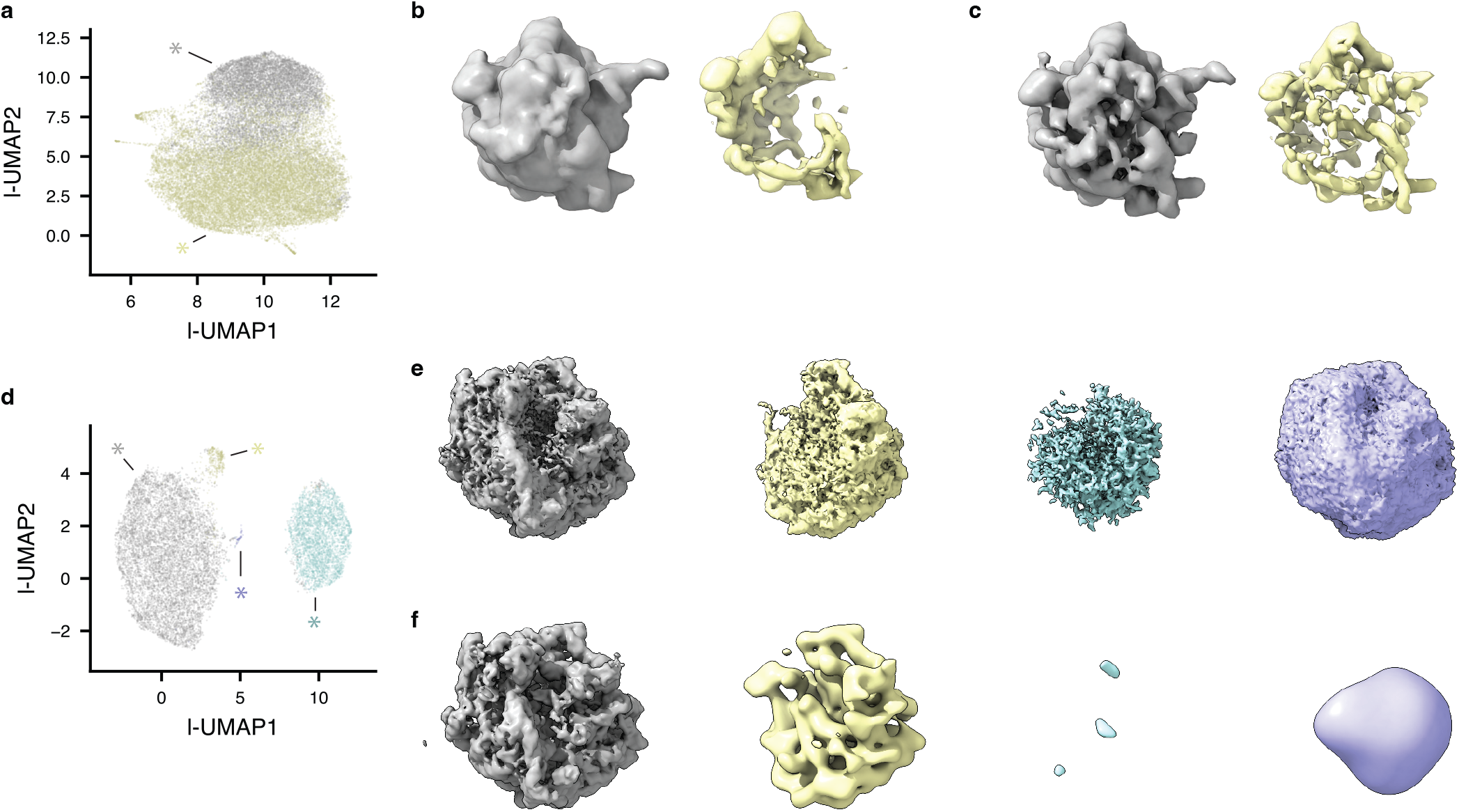
TomoDRGN analysis during iterative STA processing. **(a-c)** Data processing iteration 1 with 44,000 particles at 5x binning. **(a)** UMAP dimensionality reduction of the tomoDRGN learned latent space. **(b)** TomoDRGN-generated volumes sampled randomly within regions indicated in panel (a). Classes include 70S ribosomes (gray) and non-ribosomal false positives (yellow). **(c)** Weighted back-projection of particle sets identified with tomoDRGN and indicated in panel (a). **(d-f)** Data processing iteration 2 with 11,957 particles at 2x binning. **(d)** UMAP dimensionality reduction of the tomoDRGN learned latent space. **(e)** TomoDRGN-generated volumes sampled randomly within regions indicated in panel (d). Classes include 70S ribosomes (gray), 50S ribosomes (yellow), non-ribosomal false positives (blue), and platinum sputter (purple). **(f)** Weighted back-projection of particle sets identified with tomoDRGN and indicated in panel (d).

**Supplementary figure 4.**
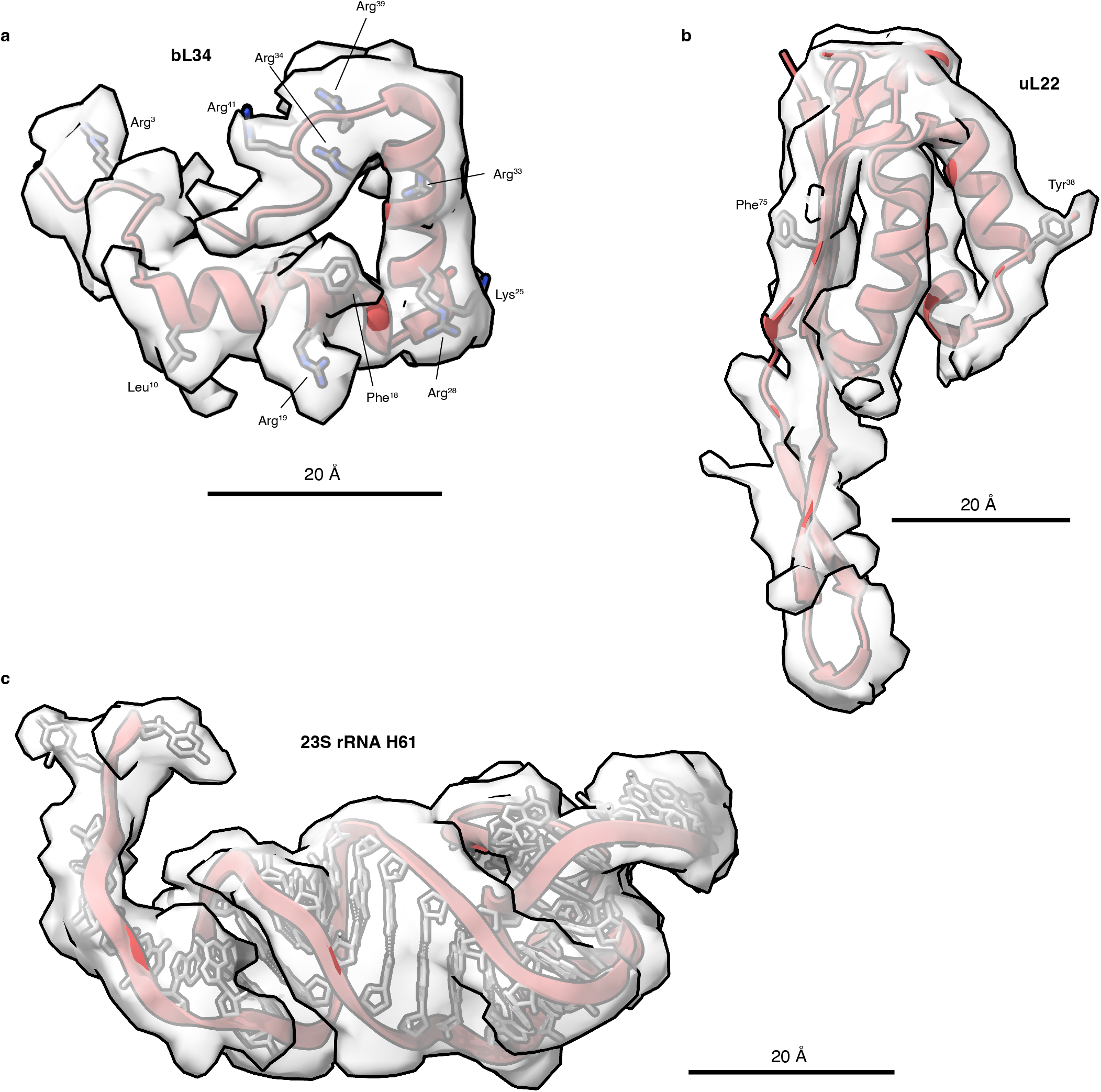
Local map quality of final 5.8 Å reconstruction. Local density (grey) of the 5.8 Å resolution Coulomb potential map (shown in Figure 2g) with atomic models (red) of ribosomal protein bL34 **(a)**, ribosomal protein uL22 **(b)**, and large subunit 23S rRNA helix 61 **(c)**. PDB model 8B0X is shown for all panels following rigid body docking into the reconstruction. Scale bar: 20 Å.

**Supplementary Table 1.** Refinement parameters used in M during round 1 of the iterative RELION-tomoDRGN-M analysis.

| Refinement | Parameters | 70S resolution (Å) |
| --- | --- | --- |
| 1 | Particle poses<br>Image warp grid 3x2 | 14.5 |
| 2 | Particle poses<br>Image warp grid 3x2<br>Stage angles | 14.4 |
| 3 | Particle poses<br>Image warp grid 3x2<br>Stage angles<br>Volume warp grid 2x3x2x10 | 14.4 |
| Notes specific to this round:<br>1. Half-maps were taken directly from RELION at 7Å/px for M species creation.<br>2. Refined tilt series containing at least 10 particles. |  |  |

**Supplementary Table 2.** Refinement parameters used in M during round 2 of the iterative RELION-tomoDRGN-M analysis.

| Refinement | Parameters | 70S resolution (Å) |
| --- | --- | --- |
| 1 | Particle poses<br>Image warp grid 6x4 | 6.0 |
| 2 | Particle poses<br>Image warp grid 6x4<br>Stage angles<br>Volume warp grid 2x3x2x10 | 5.9 |
| 3 | Particle poses<br>Image warp grid 6x4<br>Stage angles<br>Volume warp grid 2x3x2x10<br>Defocus grid search | 5.9 |
| 4 | Particle poses<br>Image warp grid 6x4<br>Stage angles<br>Volume warp grid 2x3x2x10<br>Defocus grid search<br>Tilt movie alignment | 5.9 |
| Notes specific to this round:<br>1. Half-maps were resampled in RELION from 3Å/px to 2.5Å/px prior to M species creation.<br>2. Refined tilt series containing at least 10 particles. |  |  |

**Supplementary Table 3.** Refinement parameters used in M during round 3 of the iterative RELION-tomoDRGN-M analysis.

| Refinement | Parameters | Resolution [70S] (Å) | Resolution [50S] (Å) | Refinement | Parameters | Resolution [70S] (Å) | Resolution [50S] (Å) |
| --- | --- | --- | --- | --- | --- | --- | --- |
| 1 | Particle poses<br>Image warp grid 6x4 | 6.5 | 11.7 | 7 | Particle poses<br>Image warp grid 6x4<br>Stage angles<br>Volume warp grid 2x3x2x10 | 5.9 | 13.2 |
| 2 | Particle poses<br>Image warp grid 6x4<br>Stage angles | 6.5 | 11.8 | 8 | Particle poses<br>Image warp grid 6x4<br>Stage angles<br>Volume warp grid 2x3x2x10<br>Anisotropic magnification | 5.9 | 13.3 |
| 3 | Particle poses<br>Image warp grid 6x4<br>Stage angles | 6.4 | 12.0 | 9 | Particle poses<br>Image warp grid 6x4<br>Stage angles<br>Volume warp grid 4x6x2x10 | 5.9 | 12.4 |
| 4 | Particle poses<br>Image warp grid 6x4<br>Stage angles<br>Volume warp grid 2x3x2x10 | 6.4 | 12.0 | 10 | Particle poses<br>Image warp grid 6x4<br>Stage angles<br>Volume warp grid 4x6x2x10 | 5.8 | 12.2 |
| 5 | Particle poses<br>Image warp grid 6x4<br>Stage angles<br>Volume warp grid 2x3x2x10 | 6.4 | 13.2 | 11 | Particle poses<br>Image warp grid 6x4<br>Stage angles<br>Volume warp grid 4x6x2x10 | 5.8 | 12.2 |
| 6 | Particle poses<br>Image warp grid 6x4<br>Stage angles<br>Volume warp grid 2x3x2x10 | 5.9 | 13.3 |  |  |  |  |
| Notes specific to this round:<br>1. Half-maps were resampled in RELION from 3Å/px to 2.5Å/px prior to M species creation.<br>2. Refined tilt series containing at least 10 particles for refinements 1-5, and at least 40 particles for refinements 6-11. |  |  |  |  |  |  |  |

